# Volatile emissions from diverse estuarine bacteria share core compounds with a subset of strain-specific, low abundance compounds

**DOI:** 10.64898/2026.03.24.713875

**Authors:** Eve Galen, Kajsa Roslund, Riikka Rinnan, Lasse Riemann

**Affiliations:** Center for Volatile Interactions, Department of Biology, University of Copenhagen, Universitetsparken 15, Building 1, 2100 Copenhagen E, Denmark; Marine Biological Section, Department of Biology, University of Copenhagen, Strandpromenaden 5, 3000 Helsingør, Denmark

**Keywords:** biogenic volatile organic compounds (BVOC), marine heterotrophic bacteria, volatilome, PTR-TOF-MS, headspace analysis, Baltic Sea

## Abstract

Biogenic volatile organic compounds (BVOCs) are gases that influence atmospheric chemistry, nutrient cycling, and species interactions, yet the contribution of heterotrophic marine bacteria to marine BVOC emissions remains poorly constrained. In addition, the extent to which the volatilome is linked to bacterial phylogeny is unknown. Here, we characterize the volatilome of 16 heterotrophic bacterial strains isolated from Baltic Sea surface water, spanning Alphaproteobacteria, Gammaproteobacteria, Betaproteobacteria, Bacteroidota, and Actinomycetes. Headspace BVOCs were quantified under standardized growth conditions using Proton Transfer Reaction Time-of-Flight Mass Spectrometry (PTR-TOF-MS). A broadly overlapping bacterial volatilome was identified, with compound composition and proportional abundance similar across many strains, irrespective of phylogeny. Namely, most strains shared a core set of abundant compounds with a subset of strain-specific, low abundance compounds. Acetone accounted for more than 50% of the emissions in most volatilomes. The remaining fraction of emissions were primarily comprised of other low-molecular-weight oxygenated compounds. Interestingly, two strains demonstrated strain-specific emission patterns, significantly diverging from the group in their emission rate and compound composition. Together, these findings suggest that marine heterotrophic bacteria may contribute a broadly conserved collection of BVOCs to the ocean–atmosphere interface, highlighting their role as a widespread source of trace gases in marine ecosystems.

## 1 Introduction

Biogenic volatile organic compounds (BVOCs) are generally characterized by their size (<300Da) and high vapor pressure, which enables them to easily diffuse in water and air (Weisskopf et al. 2021). These carbon-based compounds are produced by a variety of biosynthetic pathways, including primary and secondary metabolism, fermentation, sulfur metabolism, fatty acid biosynthesis, and terpene synthesis (Schulz and Dickschat 2007; Weisskopf et al. 2021). Emission of BVOCs to the atmosphere has consequences for carbon cycling as well as regional and global climate. Namely, the oxidation of BVOCs can form secondary organic aerosols, affect the atmosphere’s oxidative capacity, and impact the lifetime of greenhouse gases (Atkinson and Arey 2003; Hallquist et al. 2009; Fu et al. 2010).

Many studies have reported the production of hydrocarbons and organic sulfur compounds by phytoplankton (Simó 2001; Shaw et al. 2010; Halsey et al. 2017; Davie-Martin et al. 2020; Rocco et al. 2021; Halsey and Giovannoni 2023a), however, less understood is the role of heterotrophic bacteria in the production of these compounds and other BVOCs. Bacteria are responsible for carbon cycling in marine waters (Azam et al. 1983; Azam 1998), however, information about the identity and abundance of the BVOCs they produce is sparce. Recently, bacterial assemblages from Japanese coastal waters were found to emit acetaldehyde, acetone, and four sulfur compounds, including dimethyl sulfide (DMS) and methanethiol (MeSH; Omori et al. 2025). A few previous studies have also confirmed the production of acetone (Nemecek-Marshall et al. 1995, 1999), DMS, and MeSH (Sun et al. 2016) by marine heterotrophic bacteria as well as methyl halides (Amachi et al. 2001; Fujimori et al. 2012). No studies have reported on the total BVOC composition, named the volatilome, produced by marine bacteria. The overall bacterial contribution to marine BVOC emissions, with the possible interactive effects of these emissions, is, therefore, poorly understood.

Marine bacterial communities harbor populations with distinct metabolisms and divergent resource preferences, assimilating and transforming organic matter, including BVOCs, in diverse manners (Bunse and Pinhassi 2017). Populations may therefore have disparate effects on the cycling of specific compounds. Still, it is unknown to what extent marine bacterial populations emit characteristic BVOCs, and whether the volatilome can be linked to community composition dynamics. This information is essential for identifying environmental drivers regulating microbial BVOC emissions and for an ecological understanding of BVOC dynamics. Furthermore, research on marine BVOCs has primarily focused on targeted compounds such as isoprene, DMS and its precursors, and select oxygenated BVOCs, like methanol, acetaldehyde, and acetone. More than 3500 microbial BVOCs have been directly measured (Kemmler et al. 2025), reflecting the vast breadth of microbial emissions, yet few studies have addressed the chemical diversity of marine microbial BVOCs (Krause et al. 2025; Salinas-García et al. 2025). Studying the volatilome of marine bacteria may lend new information to distinct pathways of bacterial metabolism and insight into the bacterial contribution to pools and fluxes of specific BVOCs in marine waters.

Decoupling community-level and individual-level emissions remains a major challenge for *in situ* BVOC measurements. Currently, community-averaged measurements likely obscure strain-specific contributions to BVOC fluxes, masking the unique attributes and traits of different species. For example, single strains may be responsible for disproportionate emission rates, or emit unique volatilomes, indicating a distinctive function in the environment or an outsized impact on biogeochemical cycling of BVOCs (Weisskopf et al. 2021). Moreover, most studies do not disentangle BVOC emissions between phytoplankton and heterotrophic bacteria, with only a few studies measuring phytoplankton emissions with and without bacteria present (Moore et al. 2020; Padaki et al. 2025). Overall, only a few studies have investigated BVOC emissions from marine bacteria alone, with all studies either focusing on a single compound or a few, phylogenetically similar strains (Nemecek-Marshall et al. 1995, 1999; Thiel et al. 2009; Groenhagen et al. 2016; Lawson et al. 2020; Chhalodia et al. 2021). To date, no studies have characterized marine bacterial volatilomes across different phylogeny or used a high-sensitivity approach optimized for targeting low-molecular-weight oxygenated compounds, such as Proton Transfer Reaction Time-of-Flight Mass Spectrometry (PTR-ToF-MS).

Here, we report the emission rates and volatilomes of 16 diverse bacterial strains isolated from the Baltic Sea. Specifically, we examined the patterns between strain phylogeny and their volatilomes. As nutrient conditions, genomic variation, growth phase, and proximity to other species are known to influence the abundance and blend of BVOC emissions, all factors except genomic variation were kept constant (Choudoir et al. 2019). By comparing volatilomes across closely and distantly related strains, we aimed to study whether emission patterns are conserved along evolutionary lineages or if volatilomes reflect strain-specific adaptations.

## 2 Materials and Methods

### 2.1 Bacterial strains

Bacterial isolates were originally obtained from surface water samples (3 m) collected at the Landsort Deep station (BY31; 58° 35’ N, 18° 14’ E) in the Baltic Sea Proper between March 2003 and May 2004. Detailed procedures for water sampling, bacterial isolation, and 16S rRNA amplicon sequencing are described in Riemann et al. (2008). For the present study, 16 strains were selected from within Alphaproteobacteria, Gammaproteobacteria, Betaproteobacteria, Bacteroidota, and Actinomycetota (Table S1). Strains of varying phylogenetic relatedness were chosen for comparison, including a set of identical (in morphology and nucleotide identity) strains from different stocks for the purpose of validating our approach.

### 2.2 Bacterial cultivation and growth

Zobell marine medium of salinity 16–17 was prepared for broth and solid applications by mixing 5 g peptone 1 g yeast extract and 800 mL surface seawater from Øresund, Denmark and 200 mL Milli-Q water, and then autoclaving (Zobell 1941). M9 minimal medium (salinity 16–17) was prepared by adding 1 g NH_4_Cl, 6 g Na_2_HPO_4_*7H_2_O, 3 g KH_2_PO_4_ and 0.5 g NaCl to 1 L of Milli-Q water, autoclaved, and then adding filter sterilized MgSO_4_ and glucose to final concentrations of 2 and 22 mM, respectively (Miller 1972).

Strains were grown on Zobell agar plates. Single colonies were inoculated into Zobell broth and grown at 19L°C while shaking (250 rpm) until visible turbidity was achieved. Cells were harvested by centrifugation at 4,000 × g for 6 min. The pellet was gently resuspended in 1 mL of M9 medium. This washing step was repeated once, and each of the triplicate pellets was resuspended in 26 mL of M9 medium in 50 mL Falcon tubes. Medium-only controls were prepared in parallel.

Cell growth was monitored by optical density (OD_600_) in an Eppendorf BioPhotometer (Hamburg, Germany). After a few days of growth, cells were diluted to an OD of 0.1 in 30 mL of M9 medium in 100 mL Duran bottles. After dilution, OD was recorded every 6–12 h to identify the mid-exponential phase of growth, at which time point BVOCs were sampled from a parallel set of cultures. Immediately prior to BVOC analysis, aliquots were removed from the culture for OD measurement for growth phase confirmation and cell enumeration (fixed with 1% glutaraldehyde for 10 min at room temperature and stored at −80L°C). Cells were enumerated on a FACSCanto II flow cytometer (BD, New Jersey, USA) using SYBR Green I staining (Brussaard et al. 2010) and TrueCount beads (BD) were used to measure the flow rate.

### 2.3 Measurement of BVOCs

Duran bottles (100 mL) containing 30 mL of culture were fitted with modified lids containing an inlet and an outlet port, and incubated in a dark thermal chamber at 19L°C with shaking (250 rpm). The inlet port was connected to a zero-air generator delivering a continuous BVOC-free air flow of 4.8 L h^-1^. The outlet was connected to the PTR-ToF-MS (1000 ultra, IONICON Analytik, Austria) for real-time BVOC analysis. BVOC ion counts were monitored until a steady-state signal was reached, defined as a stable ion count for a minimum of 5 min. The time to reach steady-state varied between samples (30 - 120 min). Medium-only controls were measured daily under the same conditions as the bacterial cultures.

For measurement with the PTR-ToF-MS, hydronium (H_3_O^+^) was used as the reagent ion, and measurements were performed from a mass-to-charge ratio (*m/z*) 17 to 239 with a 0.25 Hz measurement frequency. The PTR-ToF-MS operational parameters were as follows: drift tube pressure of 2.20 mbar, temperature of 60 °C, and voltage of 500 V, corresponding to a reduced electric field (*E/N*) of 116 Td. Ion source current of 3.0 mA and a H_2_O flow of 6.0 standard cubic centimeters per minute (sccm). Instrument inlet flow was 25 sccm and temperature was 70 °C. For mass scale calibration, an internal standard (1,3-diiodobenzene, *m/z* 203.94) and the H_3_(16)O^+^ (*m/z* 21.02) peaks were used. Transmission curves and instrument calibration were achieved with a standard gas mixture (Air Liquide, Ballerup, Denmark) with known concentrations of 11 BVOCs in pure nitrogen (Table S2). The calibration data was used for compound identification. Compounds missing an authentic standard were tentatively identified using the GLOVOCS database (Yáñez-Serrano et al. 2021) in combination with a targeted literature review. Compound assignment was carried out for strain-specific *m/z* and the ten most abundant *m/z* for each strain.

### 2.4 Post-processing and BVOC emission rate calculation

Raw data from the PTR-ToF-MS were processed using PTRwid (version 3, 2018; Holzinger 2015), which performs automated mass axis calibration and peak detection, and constructs a unified mass list across samples. Peak identification and subsequent unified mass list were manually confirmed by cross-comparison with raw spectra using PTR-MS Viewer v3.4.4.13 (IONICON Analytik, Austria). Ion source-derived species (e.g., O_2_^+^, H_3_O^+^, NO_2_^+^) and interfering signals such as hydrate clusters and other artefacts were excluded from the final mass list.

Data processing, visualization, phylogenetic analysis, and statistical analysis were performed using R version 4.3.3. Raw ion counts were converted to volume mixing ratios (in parts per billion by volume (ppbv) via Equation 1.

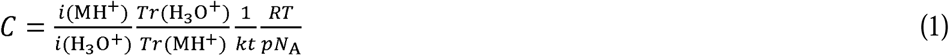

where *i*(MH^+^) is the ion current (in ion counts per second) measured for the protonated sample molecule, and i(H_3_O^+^) for the precursor ion. *N_A_* is the Avogadro’s constant, *R* is the gas constant, and *p* and *T* are the drift tube pressure and temperature, respectively, *k* is the rate coefficient, and *t* is the reaction time. Transmission factors (*Tr*) for different sample ions and the precursor ion were measured with the calibration gas mixture, to account for mass-dependence of the proton-transfer-reaction rate. Volume mixing ratio (*C*) is the partial volume of a compound in a mixture divided by the total volume, expressed here as ppbv (10^-9^ = nmol mol^-1^).

For each sample, data from the final 5 min of measurement, corresponding to the steady-state period, were averaged as the mean ppbv value for each *m/z*. Only *m/z* present in all biological replicates of a given strain were retained for analysis. Emission rates for cultures and controls were calculated following Equation 2:

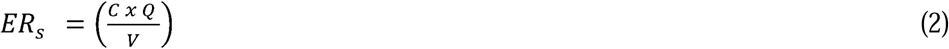

where *ER_s_*is the emission rate of the given BVOC (nmol h^-1^) in the sample; *C* is the volume mixing ratio of a VOC in the sampled air (ppbv); *Q* is the molar flow rate through the jar (mol h^-1^); and *V* is the volume (mL) of the sample.

Final emission rates were calculated following Equation 3:

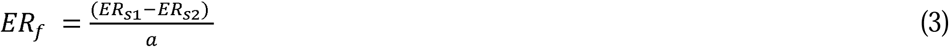

where *ER_f_* is the final emission rate of the given BVOC (nmol cell^-1^ h^-1^); *ER_s1_* is the emission rate of the BVOC in the culture; *ER_s2_* is the emission rate of the BVOC in the control; and *a* is the cell abundance (cells L^-1^). Subtraction of the control was done using a control measurement corresponding to the sample by sampling date.

### 2.5 Phylogenetic analysis

Nucleotide sequences (Table S1) were retrieved from GenBank (Benson et al. 2018) in R and aligned using the “msa” package with default parameters (Bodenhofer et al. 2015). Pairwise genetic distances were calculated from the alignment using the function “dist.alignment” in “seqinr” package (Charif and Lobry 2007) and a distance-based phylogeny was then inferred using the BIONJ algorithm implemented in the “bionj” function from the “ape” package (Paradis and Schliep 2019). Pairwise nucleotide identities were subsequently calculated from the same alignment by converting sequences to DNAbin format and computing raw distances with the “dist.dna” function in the “ape” package, which were then transformed to percent identity (1 – distance).

### 2.6 Statistical analysis

All data sets were first assessed for normality using quantile-quantile (Q-Q plots) and the Shapiro-Wilk test. Due to non-normality, the overall significance of differences in emission rates among strains and phylum/class were assessed using a Kruskal-Wallis test. The number of *m/z* emitted by each strain was normally distributed and therefore a one-way analysis of variance (ANOVA) was used to compare strains. The relationship between phylogeny and volatilome was analyzed with a principal component analysis (PCA) using SIMCA 17 (Sartorius Stedim Data Analytics AB 2021). The distance matrix was based on log-transformed and unit variance-scaled data. To assess whether the volatilomes differed between phyla and strains, the extracted principal components (PCs) were analyzed using a linear mixed-effect model (LMM) fitted with maximum likelihood (ML) using the ‘lmer’ function from the lme4 package (Bates et al. 2015). To further examine whether phylogeny may be linked with highly abundant verses lower-abundance compounds, separate PCAs were conducted on the 10 most abundant *m/z* per strain as well as the remaining *m/z*s, excluding those identified in the first analysis. Both analyses were processed following the approach above.

## 3 Results

The 16 bacterial strains belonged to Alphaproteobacteria, Gammaproteobacteria, Betaproteobacteria, Bacteroidota, and Actinomycetes (Figure 1A). Strains showed pair-wise nucleotide similarities ranging from 64 to 92%, except for isolates BAL228 and BAL217, which shared 100% similarity.

**Figure 1.**
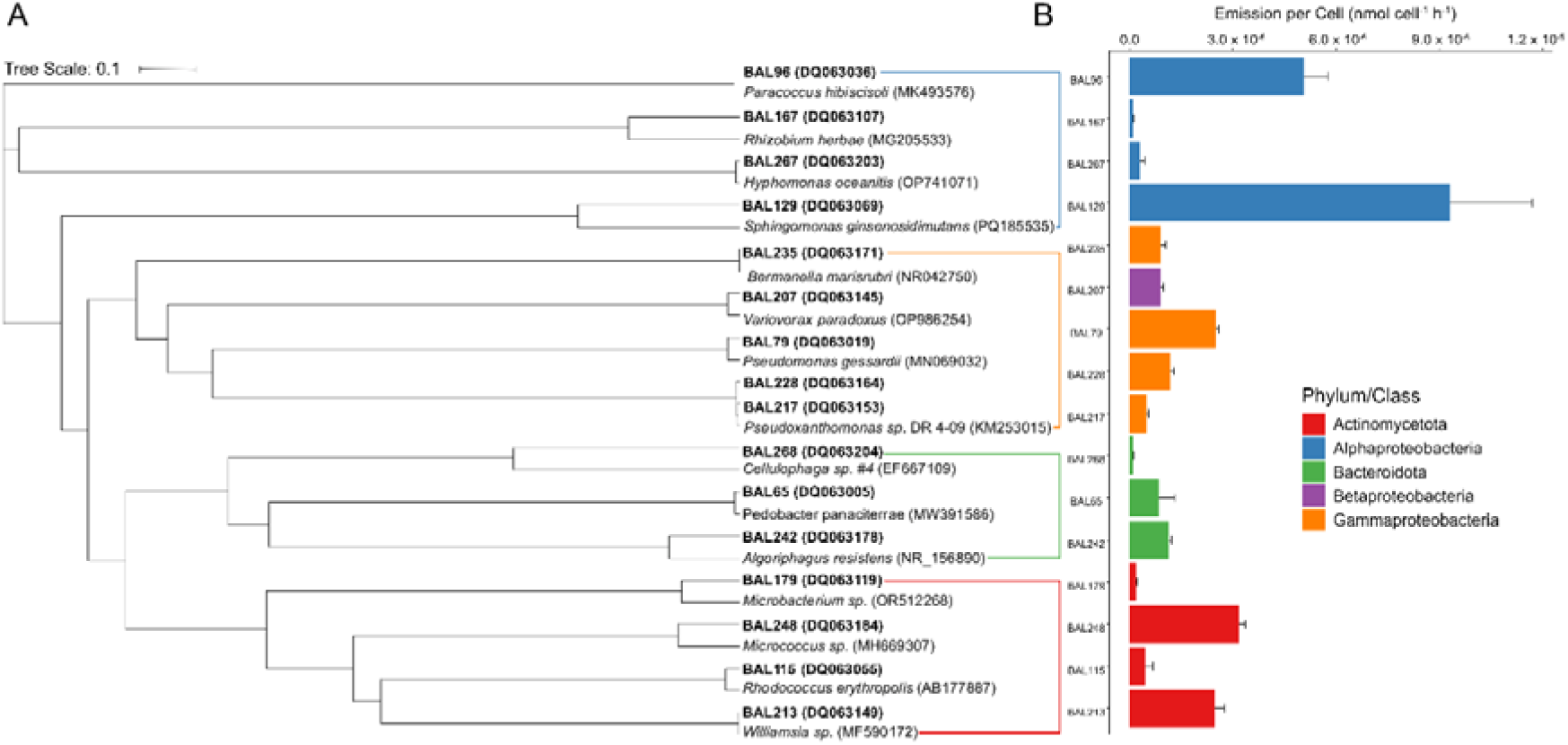
Phylogenetic tree of strains and associated total BVOC emissions. A) neighbor-joining phylogenetic tree of the 16 strains (BAL) and the nearest identified neighbor for each (Table S1). The tree scale corresponds to a 10% genetic difference between strains. B) BVOC emission rate (mean + SE, n =3 except BAL79 = 2) for each strain. Brackets and bars are colored according to phylum/class.

Quantification of BVOC emissions identified 268 unique *m/z* across all strains. Significant variation in total BVOC emission rates (nmol cell^-1^ h^-1^) was observed among strains (*p*<0.001; Kruskal-Wallis, Figure 1B). For example, BAL129 exhibited the highest BVOC emission rates (4.7 ± 1.2 × 10□ □ nmol cell^-1^ h□¹), followed by BAL96 (2.5 ± 0.4 × 10□ □ nmol cell^-1^ h□¹), while strains BAL167 and BAL268 showed the lowest emissions (5.5 ± 0.5 × 10^□8^ and 5.6 ± 0.5 × 10^□8^ nmol cell^-1^ h□¹, respectively). No differences in total emission rates were observed when strains were grouped by their phylum/class (*p*>0.5; Kruskal-Wallis).

Next, we analyzed the relative abundances of the ten most abundant *m/z* for each strain (Figure 2). These comprised 48 discrete *m/z*. Nine were tentatively matched to standards—methanol (*m/z* 33.023), an alkenyl fragment (*m/z* 41.040), acetonitrile (*m/z* 42.013), acetaldehyde (*m/z* 45.035), ethanol (*m/z* 47.051), acetone (*m/z* 59.051), isoprene (*m/z* 69.071), butanone (*m/z* 73.064) and toluene (*m/z* 93.061)—which together with the 39 other tentative compounds (Table S3), accounted for almost 90% of the BVOC profile. Notably, the “Other” fraction in BAL213 accounted for 38.7% (SD = 1.4%). Acetone was the most abundant *m/z* across most strains, comprising between 30.5 to 76.2% of the total abundance, except for BAL129, BAL79, BAL65, and BAL213. BAL129 and BAL79 exhibited the highest proportional contributions from acetaldehyde and ethanol. For BAL129, acetaldehyde accounted for 36.5% (SD = 3.0%) and ethanol 37.2% (SD = 2.2%). BAL65 was characterized by the greatest relative contribution of acetaldehyde at 64.6% (SD = 1.6%). In contrast, BAL213 showed a spread across multiple lower-abundance *m/z* signals, with *m/z* 87.078 and *m/z* 101.092 accounting for the highest proportional abundances at 12.0% (SD = 9.3%) and 12.0% (SD = 1.2%), respectively. *m/z* 31.012 (formaldehyde), methanol, and an alkenyl fragment were also prevalent across strains. Formaldehyde accounted for an average of 3.2% (SD = 3.0%) of relative emissions, methanol for 6.9% (SD = 4.9%), and the alkenyl fragment for 5.9% (SD = 6.7%). The emission rate per hour per cell for formaldehyde, methanol, the alkenyl fragment, acetaldehyde, ethanol, acetone, *m/z* 87.078, and *m/z* 101.092 is presented in Table S4.

**Figure 2.**
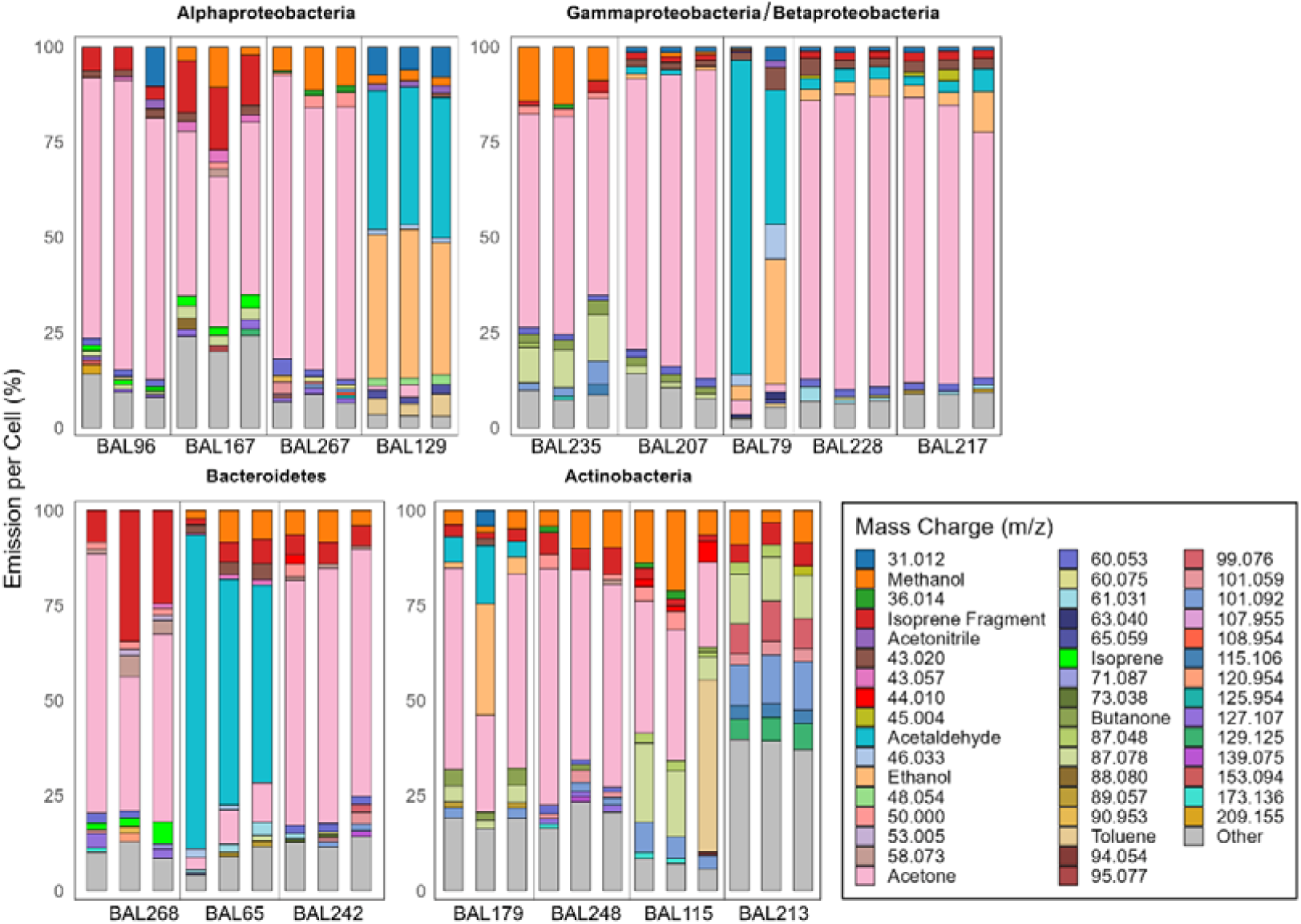
Relative abundance of BVOCs (% of total) for each replicate strain. Strains are grouped by Phylum/Class. The ten most abundant *m/z* for each strain are included, and bars are stacked from top to bottom in *m/z* order, with the grey proportion at the bottom, called “Other”, representing the sum of the rest of the masses emitted. Known *m/z*, tentatively identified using standards, are indicated by their chemical name in the figure legend.

Between 42 and 182 distinct *m/z* were detected across strains (Table 1). The number of *m/z* produced by each strain was not related to the phylogeny of the strains (*p*>0.5; ANOVA). Eight strains (BAL96, BAL267, BAL129, BAL207, BAL228, BAL217, BAL248, BAL213) emitted strain-specific *m/z* (Table 2). These strain-specific *m/z* were in the lower 0.01% of all emissions, with the most unique *m/z* accounting for less than 0.001% of total emissions. Only the *m/z* 83.085 (C_6_H_10_H^+^) ranked in the top 100 emitted compounds (top 0.03% of all emissions). It accounted for 0.9% of emissions from BAL213 and was the 21^st^ most abundant compound emitted by this strain.

**Table 1.** Total number of mass-to-charge ratios (*m/z*) and unique *m/z* for each bacterial strain. Strains are ordered by phylogeny.

|  | Alphaproteobacteria |  |  |  | Gammaproteobacteria/Betaproteobacteria |  |  |  |  | Bacteriodota |  |  | Actinomycetota |  |  |  |
| --- | --- | --- | --- | --- | --- | --- | --- | --- | --- | --- | --- | --- | --- | --- | --- | --- |
| BAL | 96 | 167 | 267 | 129 | 235 | 207 | 79 | 228 | 217 | 268 | 65 | 242 | 179 | 248 | 115 | 213 |
| Total $m/z$ | 133 | 116 | 80 | 71 | 53 | 127 | 104 | 182 | 136 | 77 | 107 | 114 | 111 | 134 | 42 | 158 |
| Unique $m/z$ | 2 | 0 | 1 | 12 | 0 | 1 | 0 | 1 | 1 | 0 | 0 | 0 | 0 | 5 | 0 | 4 |

**Table 2.**
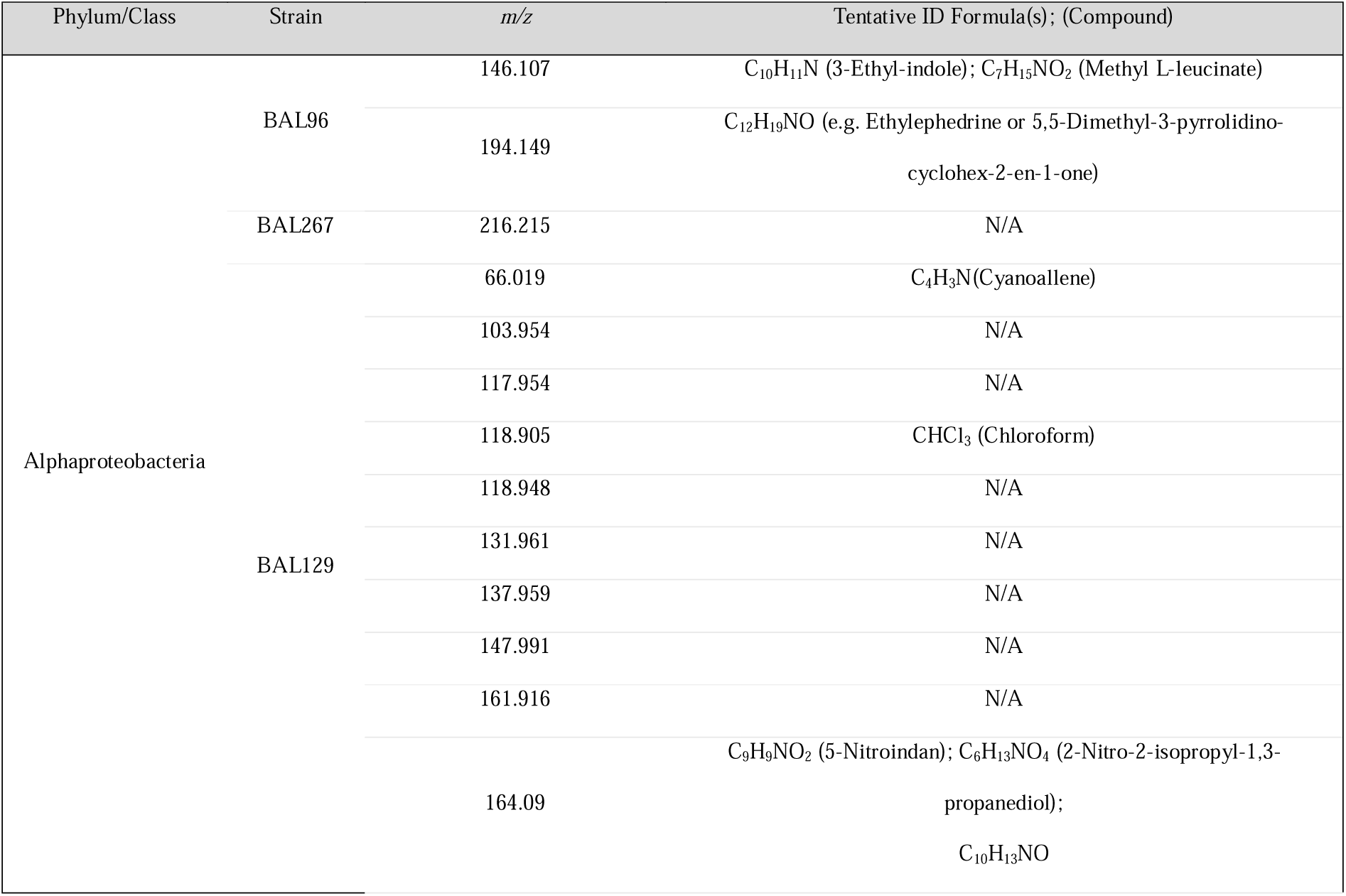
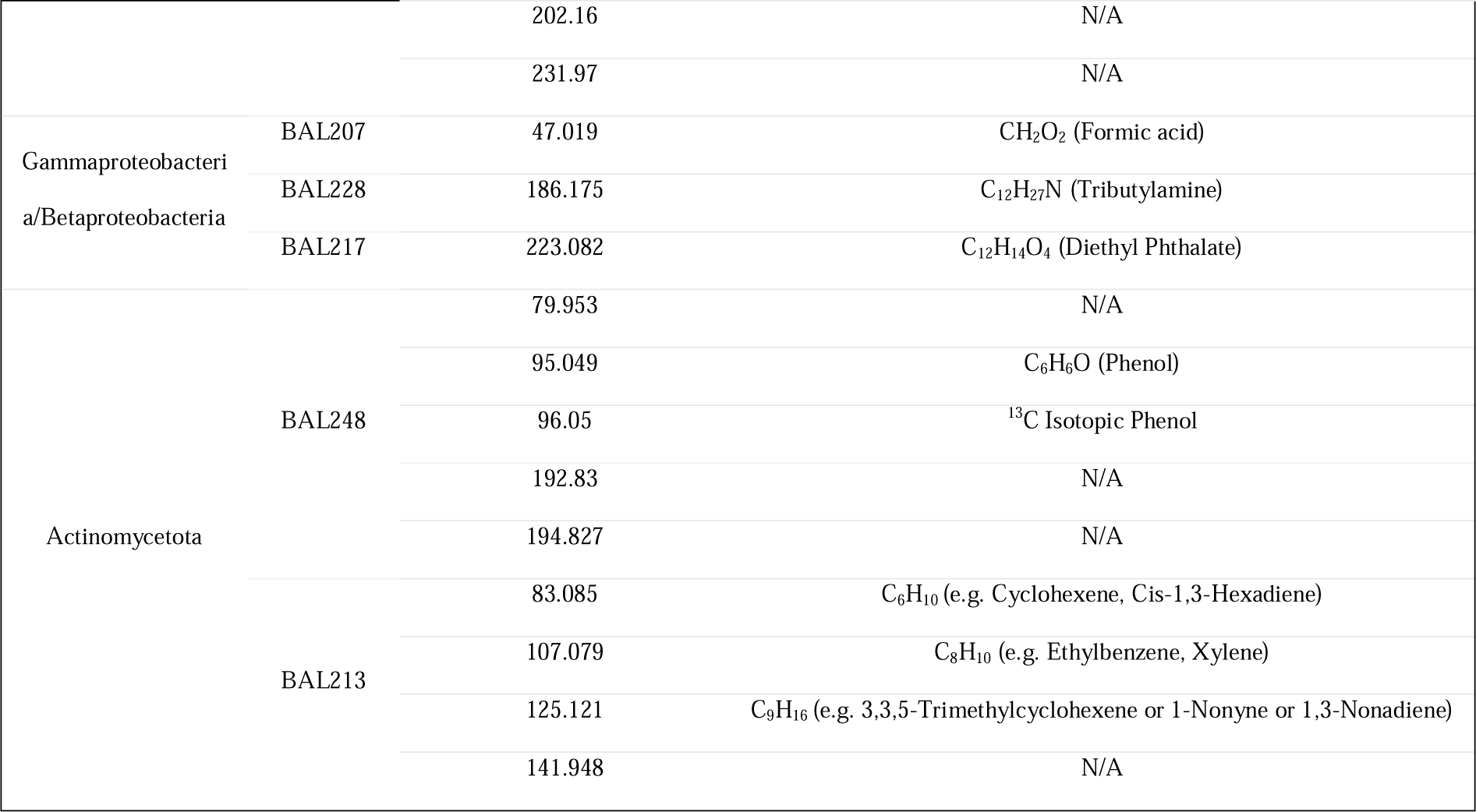
Strain-specific mass-to-charge ratios (*m/z*) and their tentative identification. N/A indicates no identification was found. Tentative identification was carried out using GLOVOCS database (Yáñez-Serrano et al. 2021) in combination with a targeted literature review.

The relationship between phylogeny and volatilome was analyzed using a PCA (Figure 3). The first two PCs accounted for 42.0% of the total variance. The strains did not cluster based on their phylogeny (*p*>0.5; LMMs on the PC scores). Notably, Bacteroidota (BAL65, BAL242, BAL268) and Gammaproteobacteria (BAL79, BAL217, BAL228, BAL235) occupied the center of the score plot, alongside several Alphaproteobacteria and Actinomycetota strains, suggesting a similar volatilome among these groups. In contrast, BAL129 (Alphaproteobacterium) and BAL213 (Actinomycetota) were divergent on the plot, indicating distinctive volatilomes.

**Figure 3.**
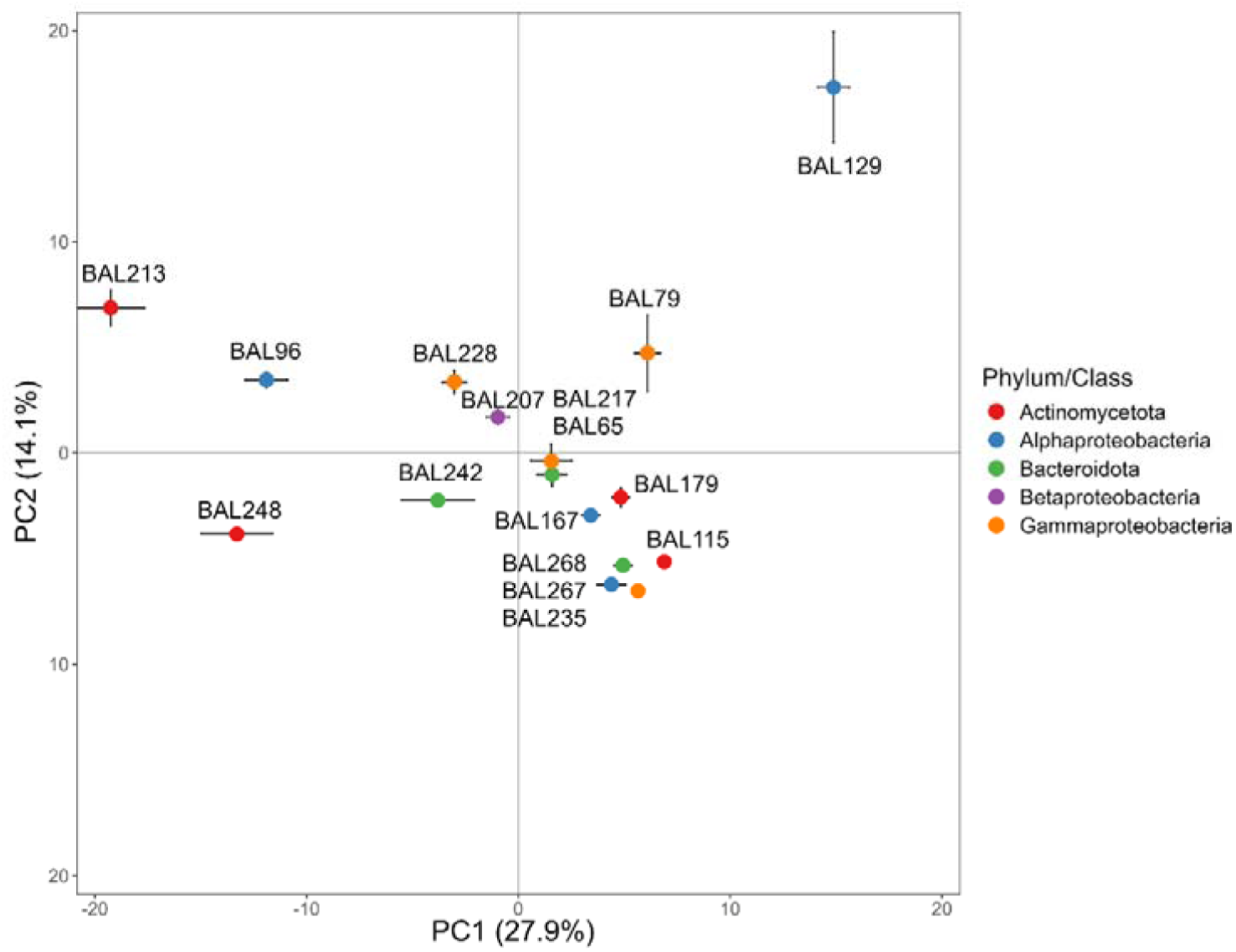
Principal component analysis (PCA) score plot of the total volatilome for the 16 strains (mean PC score position ± SE, n=3 except BAL79 = 2). The volatilomes encompassed 268 unique *m/z*.

Two additional PCAs were conducted to determine whether strain-level differences in volatilome structure were driven primarily by the most abundant compounds or by the lower-abundance fraction. First, a PCA was performed on the 10 most abundant compounds per strain (48 distinct *m/z*; corresponding to colored fraction in Figure 2), explaining 28.6% of the variance in the first two PCs (Figure S1A). This analysis showed that strains clustered tightly near the center of the score plot, indicating high similarity in the emission blend across strains. Second, a PCA was performed on the remaining volatilome after removal of the 10 most abundant compounds per strain (220 distinct *m/z*; corresponding to the “Other” fraction in Figure 2), which explained 38.5% of the variance in the first two PCs (Figure S1B). Here, strains spread across the score plot in a similar pattern to the whole volatilome PCA (Figure 3), indicating that much of the strain-level variance was captured by the lower-abundance compounds rather than the dominant compounds.

A pairwise comparison of shared *m/z* among strains revealed a few cases where phylogenetic relatedness and volatilome similarity appear linked (Figure 4). For instance, the identical Gammaproteobacteria BAL228 and BAL217 shared the greatest number of *m/z* (127). Certain strains exhibited a consistently low degree of overlap across the entire panel, regardless of phylogenetic affiliation. For example, BAL129, BAL235, and BAL115 shared relatively few *m/z* features with all other strains. In contrast, BAL213, BAL96, BAL167, and BAL217 shared a relatively high number of features with multiple other strains.

**Figure 4.**
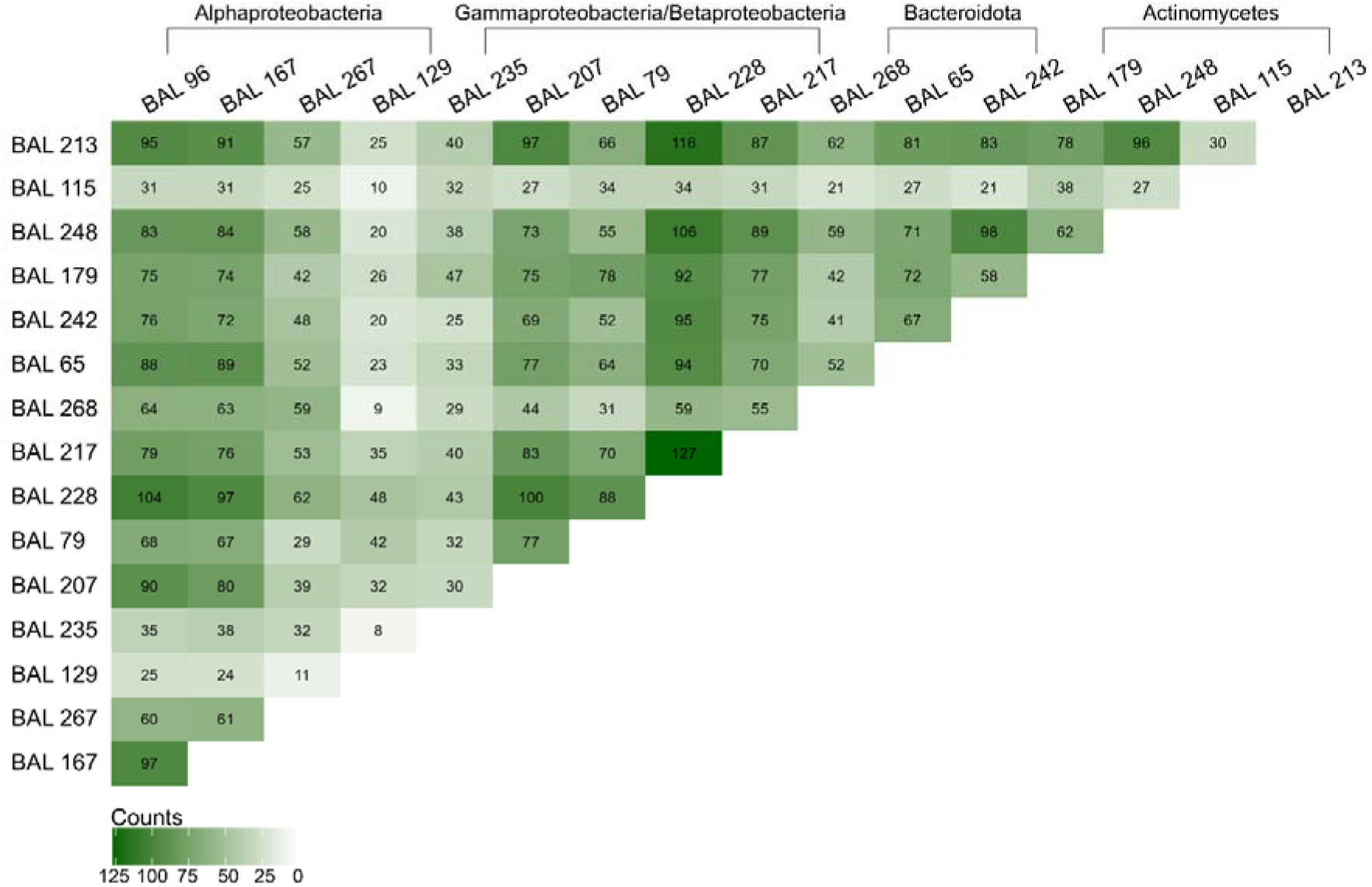
Heatmap showing the total number of mass-to-charge ratio (*m/z*) each strain shares with each other. Dark green indicates more shared *m/z*, while light green indicates less shared *m/z*. The number of shared *m/z* is shown inside each rectangle. Strains are ordered by phylogeny.

## 4 Discussion

In this study we show that 16 phylogenetically diverse Baltic Sea strains emit a rich blend of BVOCs, including both a shared core set of abundant compounds and a subset of strain-specific, low-abundance compounds. The emission patterns did not reflect phylogeny, apart from those strains that shared identity. Overall, this indicates that evolutionary relatedness was not a primary driver of the volatilome structure. Rather, our results suggest the presence of a conserved bacterial volatilome shared across taxa, complemented by a minor subset of strains with distinct emission profiles.

### 4.1 Key compounds in the bacterial volatilome

Across the 16 strains, acetone represented the largest fraction of emissions (50.2%). Similarly, acetone emissions from marine bacteria have been described as early as 1995 by Nemecek-Marshall et al., who observed substantial production in *Vibrio* isolates from coastal waters off California. Recently, Omori et al. (2025) reported acetone concentrations ∼10x higher than MeSH and ∼100x higher than acetaldehyde in association with bacterial assemblages from Japanese coastal waters. In both studies, acetone increased with growth and was highest towards the stationary phase, indicating that acetone emissions from the strains measured here could increase in later growth stages. Importantly, acetone production was not universal across the analyzed *Vibrio* isolates (Nemecek-Marshall et al. 1995), consistent with our observation that acetone is a dominant bacterial BVOC but not ubiquitous. Together, the high acetone emission rates observed both in this study, and in others, suggests that acetone is a key compound emitted by marine bacteria. Moreover, the differences in acetone production between bacterial strains suggest that shifts in bacterial abundance and composition could impact acetone fluxes to the atmosphere. Once in the climate system, acetone can enhance ozone formation and alter the lifetime of methane, acting as a precursor to changes in net radiative forcing (Folkins and Chatfield 2000; Wang et al. 2020). Collectively, these observations suggest that bacterial acetone production should be considered in oceanic acetone cycling models and in the climate system.

Other compounds measured in relatively high abundance included acetaldehyde, ethanol, and methanol. These low-molecular-weight oxygenated compounds have been widely observed in marine ecosystems (Sinha et al. 2007; Beale et al. 2013; Dixon et al. 2013; Halsey et al. 2017; Davie-Martin et al. 2020; Zhou et al. 2023) and can originate from bacteria (Kuzma et al. 1995; Bunge et al. 2008; Roslund et al. 2021; Salinas-García et al. 2026). Yet in the marine environment, few studies have quantified their emission directly from heterotrophic bacteria (Nemecek-Marshall et al. 1995, 1999; Thiel et al. 2009; Groenhagen et al. 2016; Lawson et al. 2020; Chhalodia et al. 2021), with production commonly attributed to phytoplankton, and the role of bacteria commonly described solely in terms of consumption and degradation (Halsey et al. 2017; Wohl et al. 2020; Davie-Martin et al. 2020; Halsey and Giovannoni 2023). In culture studies, acetaldehyde emissions are commonly associated with early exponential growth (Bunge et al. 2008; Omori et al. 2025; Salinas-García et al. 2026). This timing fits with our study, indicating that those strains that did not produce acetaldehyde can be considered non-emitters. For ethanol and methanol, production has been documented in clinically relevant heterotrophic bacteria (Bunge et al. 2008; McNerney et al. 2012; Sovová et al. 2013; Chippendale et al. 2014), but, to our knowledge, no comparable observations exist for marine heterotrophic bacteria. It is important to note that no sulfur compounds, including DMS or MeSH, were tentatively identified in our study despite their ubiquity in the marine environment and known origin from marine bacteria (Brock et al. 2014; Omori et al. 2025). This may reflect both the use of minimal medium and the timing of the measurements. Production of sulfur compounds by heterotrophic bacteria is often linked to the availability of sulfur-containing precursors, such as amino acids or dimethylsulfoniopropionate (DMSP), which were not present in our minimal media (Brock et al. 2014; Roslund et al. 2021; Salinas-García et al. 2026). Likewise, sulfur emissions are often observed during the stationary to death phase when cells have produced complex exudates and cell lysis has deposited organic precursors into the medium for late stage bacteria to utilize (Baumler et al. 2007; Nawrath et al. 2012). Overall, these results highlight that even the emission of the most abundant compounds are highly linked with growth stage and nutrient availability.

### 4.2 A conserved bacterial volatilome

The analysis identified a broadly overlapping bacterial volatilome during the active growth phase, with compound composition and proportional abundance similar across most strains, irrespective of phylogeny. Gammaproteobacteria, Betaproteobacteria, and Bacteroidota had relatively conserved volatilomes, while Actinomycetota and Alphaproteobacteria included strains showing larger variability. This observation is consistent over the total volatilome (Figure 3), the blend of most abundant compounds (Figure S1A), and the blend of lower-abundance compounds (Figure S1B).

Of the *m/z* observed, between 21 and 116 *m/z* (average 58 *m/z*) were shared across strains of different phylogeny, supporting the presence of a broadly overlapping volatilome (Figure 4). This excludes BAL129, which shared comparatively few (average 24 *m/z*) with the other strains. No single *m/z* was emitted by all strains, although most strains emitted large amounts of acetone (Figure 2). Overall, the shared bacterial volatilome appears to be dominated by oxygenated BVOCs. Prior work across various ecosystems has observed a similar pattern of emission from heterotrophic bacteria. For example, culture-based studies have shown that diverse bacterial taxa produce a shared set of low-molecular-weight BVOCs despite differences in phylogeny (Bos et al. 2013; Fitzgerald et al. 2021). These commonly shared compounds are often small oxygenated molecules such as alcohols, ketones, and aldehydes, that arise from conserved pathways of central metabolism (Schulz and Dickschat 2007; Weisskopf et al. 2021). Additionally, carbon substrate composition impacts volatilome composition (Rath et al. 2018; Timm et al. 2018; Jenkins and Bean 2020; Rajendran et al. 2024). Since glucose was the sole carbon source, our emissions likely originate from glycolysis and downstream pathways centered on pyruvate and acetyl-CoA (Filipiak et al. 2012). Acetone, acetaldehyde, ethanol, and other low-molecular-weight oxygenated compounds are common end-products of these pathways (Filipiak et al. 2012; Fitzgerald et al. 2021). This suggests that the overlap observed here reflects emissions of common metabolic end-products from active growth rather than phylogenetically restricted pathways.

### 4.3 Strain-level differentiation of volatilomes

The conserved volatilome observed in our study was not universal, as BAL129 (Alphaproteobacteria) and BAL213 (Actinomycetes) clearly diverged from the other strains. BAL129 had the highest emission rate of all strains (Figure 1B), the lowest number of shared *m/z* (Figure 4), and the highest number of unique *m/z* (Table 1). Interestingly, its volatilome was dominated by acetaldehyde and ethanol, with little acetone and no alkenyl fragment production (Figure 2). BAL129 is identical at the 16S rRNA nucleotide level to the soil bacterium *Sphingomonas ginsenosidimutans* (Choi et al. 2010). To our knowledge, BVOC emissions from *S. ginsenosidimutans* have not been examined. Within the *Sphingomonas* genus, BVOC emissions have only been documented from *Pseudomonas sp.* strain 19-rlim and *Sphingomonas sp.* strain BIR2-rlima for the terpene 2-methylisoborneol, an odorous compound sometimes found in drinking water (Eaton 2012). It is unknown whether the high BVOC emission found in BAL129 is a general feature in *Sphingomonas*, or specific to our strain. Given that *Sphingomonas* are widely distributed in marine environments (Eguchi et al. 1996; Cavicchioli et al. 1999; Yang et al. 2020), they have the potential to affect overall marine BVOC emissions, and a future volatilomic analysis of this genus would be warranted.

BAL213 also differed considerably from the other strains, particularly because its volatilome was dominated by less abundant compounds (Figure 2). In addition, BAL213 did not emit acetone, acetaldehyde, or ethanol. Rather, it produced compounds with higher *m/z*, such as 87.078 (C_5_H_10_O), 99.076 (C_6_H_10_O), and 101.092 (C_6_H_12_O). BAL213 is identical at the 16S rRNA nucleotide level to the bacterium *Williamsia sp. strain A155,* isolated from Atacama Desert soil (accession MF590172). In a previous study, the strain W*illiamsia sp.* FLCC425 was shown to emit a distinct volatilome, and to produce relatively few BVOCs compared to the other Actinobacteria measured (Choudoir et al. (2019). The distinctive volatilome observed for this genus may indicate specialized physiology. In marine waters where low-abundance taxa contribute outsized functional effects, *Williamsia* could uniquely impact the pools and cycling of low-abundance reactive gases.

## 5 Conclusions

Using real-time PTR-ToF-MS volatilomics, we show that phylogenetically diverse heterotrophic bacteria from Baltic Sea surface waters emit chemically complex BVOC blends that include a shared core volatilome with strain-specific differentiation and no clear link with phylogeny. Our findings support a trait-based view in which strain-level physiology and substrate availability contribute more strongly to volatilome structure than phylogenetic relatedness (Audrain et al. 2015; Schulz-Bohm et al. 2017; Meredith and Tfaily 2022). This has implications for how bacterial contributions are represented in marine BVOC budgets, as heterotrophic bacterial emissions are still largely ignored when modeling marine BVOC cycling. Next steps should expand beyond exponential growth and single-substrate conditions by testing across growth-phases, with diverse carbon sources, and with mixed communities where interactions can be studied. In parallel, pairing volatilomics with transcriptomics and metabolomics is key for attributing BVOC production to bacterial metabolism. Nevertheless, our results demonstrate that heterotrophic marine bacteria emit strain-specific volatilomes with shared core compounds, supporting the need to incorporate both these abundant compounds and bacterial trait variation into marine microbial models, BVOC budgets, and flux estimates.

## Author contributions

**Eve Galen:** Conceptualization, Investigation, Formal analysis, Writing – original draft, Writing – review & editing. **Kajsa Roslund:** Investigation (instrument operation), Writing – review & editing. **Riikka Rinnan:** Funding acquisition, Writing – review & editing. **Lasse Riemann:** Conceptualization, Funding acquisition, Writing – review & editing.

## Acknowledgements

We thank Cecilie Fryland Appeldorff for assistance with flow cytometry, Joseph Donald Martin for help with laboratory work, and Tamás Plaszkó for advice on data processing and analysis. We thank the Danish National Research Foundation (Center for Volatile Interactions – VOLT, DNRF168) for funding this research.

## Data Availability Statement

The data that supports the findings of this study are available at https://www.erda.dk/archives/0eb44b6c3d2a5fdac1e549d7694985c3/published-archive.html.

## Supporting Information

Additional supporting information can be found online in the Supporting Information section.

