## Supporting Information for "Volatile emissions from diverse estuarine bacteria share core compounds with a subset of strain-specific, low abundance compounds"

**Table S1.** Taxonomic identification for the used isolates and their nearest neighbors in GenBank.

| **Measured Strains** | | | | | **Nearest Neighbor** | | | | | |
| --- | --- | --- | --- | --- | --- | --- | --- | --- | --- | --- |
| **Name** | **Accession**  **number** | | **Strain** | **Phylum** | **Nearest neighbor** | **Accession**  **number** | **Query cover (%)** | **Percent identity (%)** | **Environment** | **Reference** |
| BAL96 | | DQ063036 | Rhodobacter | Alphaproteobacteria | *Paracoccus hibiscisoli* strain H02Y-003A | MK493576 | 100 | 100.00 | Algal culture | Direct Submission |
| BAL167 | | DQ063107 | Rhizobiaceae | Alphaproteobacteria | Rhizobium herbae | MG205533 | 100 | 99.75 | Soil | Direct Submission |
| BAL267 | | DQ063203 | Hyphomonas | Alphaproteobacteria | Hyphomonas oceanitis | OP741071 | 100 | 100.00 | NA | Direct Submission |
| BAL129 | | DQ063069 | Sphingomonadales | Alphaproteobacteria | Sphingomonas ginsenosidimutans strain HNXS-W42 | PQ185535 | 99 | 100.00 | South China Sea | Direct Submission |
| BAL235 | | DQ063171 | Oceanobacter | Gammaproteobacteria | Bacterium DG940 | NR_042750 | 100 | 100.00 | Red Sea | Pinhassi and Berman 2003 |
| BAL207 | | DQ063145 | Comamonadaceae | Betaproteobacteria | Variovorax paradoxus | OP986254 | 100 | 100.00 | NA | Twizeyimana et al. 2023 |
| BAL79 | | DQ063019 | Pseudomonadaceae | Gammaproteobacteria | Pseudomonas gessardii strain ST3SE | MN069032 | 100 | 100.00 | Soil | Direct Submission |
| BAL228 | | DQ063164 | Xanthomonadaceae | Gammaproteobacteria | Pseudoxanthomonas sp DR4-09 | KM253015 | 100 | 100.00 | Root | Direct Submission |
| BAL217 | | DQ063153 | Xanthomonadaceae 2 | Gammaproteobacteria | Pseudoxanthomonas sp DR4-09 | KM253015 | 100 | 100.00 | Root | Direct Submission |
| BAL268 | | DQ063204 | Flavobacteriaceae | Flavobacteriia | Cellulophaga baltica | EF667109 | 100 | 95.69 | Sea-water | Holmfeldt et al. 2007 |
| BAL65 | | DQ063005 | Sphingobacteriaceae | Sphingobacteriia | Pedobacter panaciterrae strain SCZ9 | MW391586 | 100 | 99.09 | NA | Direct Submission |
| BAL242 | | DQ063178 | Flexibacteriaceae | Cytophagia | Algoriphagus resistens | NR_156890 | 100 | 99.54 | Marine sediment | Han et al. 2017 |
| BAL179 | | DQ063119 | Microbacteriaceae | Actinomycetes | Microbacterium sp. | OR512268 | 100 | 99.75 | Marine sediment | Direct Submission |
| BAL248 | | DQ063184 | Micrococcaceae | Actinomycetes | Micrococcus sp. Strain H390 | MH669307 | 100 | 99.53 | Pine tree | Direct Submission |
| BAL115 | | DQ063055 | Mycobacteriaceae | Actinomycetes | Rhodococcus erythropolis | AB177887 | 100 | 100.00 | NA | Futamata et al. 2004 |
| BAL213 | | DQ063149 | Mycobacteriaceae 2 | Actinomycetes | Williamsia sp. | MF590172 | 100 | 100.00 | Soil | Direct Submission |

**Table S2.** Composition of the gas mixture used for calibration and transmission curves in the Proton Transfer Reaction Time-of-Flight Mass Spectrometry.

| **Compound** | **Molecular formula** | **Molecular mass (u)** |
| --- | --- | --- |
| Acetaldehyde | C_2_H_4_O | 44.05 |
| Methanol | CH_4_O | 32.04 |
| Ethanol | C_2_H_6_O | 46.07 |
| Acetonitrile | C_2_H_3_N | 41.05 |
| Acetone | C_3_H_6_O | 58.08 |
| Isoprene | C_5_H_8_ | 68.12 |
| 2-butanone | C_4_H_8_O | 72.11 |
| Benzene | C_6_H_6_ | 78.11 |
| Toluene | C_7_H_8_ | 92.14 |
| m-Xylene | C_8_H_10_ | 106.16 |
| α-Pinene | C_10_H_16_ | 136.24 |

**Table S3.** Tentative identification of the ten most abundant mass-to-charge ratio (*m/z*) for each strain (36 *m/z* in total). N/A indicates no identification was found. The identification was carried out using the GLOVOCS database (Yáñez-Serrano et al. 2021) in combination with literature.

| ***m/z*** | **Tentative Formula** | **Tentative Compound(s)** |
| --- | --- | --- |
| 31.012 | (CH_2_O)H^+^ | Formaldehyde |
| 36.014 | N/A | N/A |
| 43.02 | (C_2_H_2_O)H^+^ | Acetyl fragment |
| 43.057 | (C_3_H_6_)H^+^ | Propene or alkyl fragment |
| 44.01 | (CHNO)H^+^ | Fulminic acid or isocyanic acid |
| 45.004 | N/A | N/A |
| 46.033 | (CH_3_NO)H^+^ | Formamide |
| 48.054 | (CH_5_NO)H^+^ | o-Methylhydroxylamine or methoxyam |
| 50.003 | N/A | N/A |
| 53.005 | N/A | N/A |
| 58.073 | (C_3_H_7_N)H^+^ | 1-Methylethenylamine or 2-propanimine |
| 60.053 | (C_3_H_7_O)H^+^ | Carbonyl fragment |
| 60.075 | (C_3_H_9_N)H^+^ | Propan-1-amine or propylamine |
| 61.031 | (C_2_H_4_O_2_)H^+^ | Acetic acid |
| 65.059 | (C_2_H_8_O_2_)H^+^ | Ethylene dihydrate |
| 71.087 | (C_5_H_10_)H^+^ | Alkyl fragment (several compounds) |
| 73.038 | (C_3_H_4_O_2_)H^+^ | 2-Propenoic acid or acrylic acid |
| 87.048 | (C_4_H_6_O_2_)H^+^ | Carboxyl-acid derivatives (several compounds) |
| 87.078 | (C_5_H_10_O)H^+^ | Pentanone, pentanal, or 2-methylbutanal |
| 88.08 | (C_4_H_9_NO)H^+^ | N,N-Dimethyl-acetamide |
| 90.953 | (C_2_H_2_S_2_)H^+^ | 1,2-Dithiete |
| 94.054 | (C_6_H_7_N)H^+^ | Aniline, methylpyridine,  or N-2-oropynyl-2-propyn-1-amine |
| 95.077 | (C_3_H_10_O_3_)H^+^ | Methanol ethylene glycol |
| 99.076 | (C_6_H_10_O)H^+^ | Hexenals (several compounds) |
| 101.059 | (C_5_H_8_O_2_)H^+^ | Acetylacetone |
| 101.092 | (C_6_H_12_O)H^+^ | Hexanal, hexanone, or cis-3-hexenol |
| 107.955 | N/A | N/A |
| 108.954 | (C_2_H_5_Br)H^+^ | Ethyl bromide |
| 115.106 | (C_7_H_14_O)H^+^ | Heptanal, heptanone,  or 2,4-dimethyl-3-pentanone |
| 125.954 | N/A | N/A |
| 127.107 | (C_8_H_14_O)H^+^ | 6-methyl-5-hepten-2-one (sulcatone)  or 1-Octen-3-one, octenal |
| 129.125 | (C_8_H_16_O)H^+^ | Octanal, octanone, 1-octen-3-ol,  or 2,2,4-trimethyl-3-pentanone |
| 139.075 | (C_8_H_10_O_2_)H^+^ | Creosol |
| 153.094 | (C_9_H_12_O_2_)H^+^ | 4-Ethyl guiacol |
| 173.136 | (C_13_H_16_)H^+^ | 3-heptynylbenzene |
| 209.155 | (C_13_H_20_O_2_)H^+^ | Carvyl propionate |

**Table S4.** Total BVOC emission rates (nmol cell^-1^ h^-1^ ± SE) of select compounds for each strain. Compound selection is based on high relative abundance and strain relevance (see Results for elaboration). N/A indicates no signal was detected. The *m/z* with associated chemical names were identified using standards.

| **Strain** | **31.012** | **Methanol**  **(33.023)** | **Alkenyl Fragment**  **(41.04)** | **Acetaldehyde**  **(45.035)** | **Ethanol**  **(47.051)** | **Acetone (59.051)** | **87.078** | **101.092** |
| --- | --- | --- | --- | --- | --- | --- | --- | --- |
| BAL96 | 7.16 ± 6.14 x 10^-8^ | N/A | 1.04 ± 0.21 x 10^-7^ | N/A | N/A | 1.39 ± 0.66 x 10^-6^ | 2.44 ± 0.41 x 10^-8^ | 1.17 ± 0.32 x 10^-8^ |
| BAL167 | N/A | 9.15 ± 3.32 x 10^-10^ | 2.56 ± 0.40 x 10^-9^ | N/A | N/A | 7.77 ± 1.56 x 10^-9^ | 5.50 ± 1.25 x 10^-10^ | 2.43 ± 0.89 x 10^-10^ |
| BAL267 | N/A | 2.06 ± 0.45 x 10^-9^ | N/A | N/A | N/A | 1.79 ± 0.68 x 10^-8^ | N/A | 1.62 ± 0.36 x 10^-10^ |
| BAL129 | 3.44 ± 1.01 x 10^-7^ | 1.09 ± 0.22 x 10^-7^ | N/A | 1.70 ± 0.44 x 10^-6^ | 1.72 ± 0.43 x 10^-6^ | 5.14 ± 2.23 x 10^-8^ | N/A | N/A |
| BAL235 | N/A | 1.18 ± 0.17 x 10^-8^ | 1.62 ± 0.83 x 10^-9^ | N/A | N/A | 5.15 ± 0.44 x 10^-8^ | 9.99 ± 1.76 x 10^-9^ | 3.33 ± 1.50 x 10^-9^ |
| BAL207 | 4.95 ± 0.46 x 10^-9^ | 3.59 ± 0.52 x 10^-9^ | 5.78 ± 0.21 x 10^-9^ | 4.50 ± 1.01 x 10^-9^ | 4.10 ± 0.43 x 10^-9^ | 2.75 ± 0.27 x 10^-7^ | 6.09 ± 0.64 x 10^-9^ | 3.32 ± 0.63 x 10^-9^ |
| BAL79 | 2.68 ± 2.03 x 10^-8^ | 6.14 ± 0.16 x 10^-9^ | 2.88 ± 0.56 x 10^-9^ | 7.34 ± 2.79 x 10^-7^ | 2.34 ± 1.88 x 10^-7^ | 3.72 ± 1.02 x 10^-8^ | 3.19 ± 0.60 x 10^-9^ | 1.32 ± 0.22 x 10^-9^ |
| BAL228 | 6.35 ± 0.62 x 10^-9^ | 7.70 ± 2.11 x 10^-10^ | 9.89 ± 1.18 x 10^-9^ | 1.67 ± 0.20 x 10^-8^ | 1.91 ± 0.46 x 10^-8^ | 3.92 ± 0.42 x 10^-7^ | 1.62 ± 0.15 x 10^-9^ | 1.20 ± 0.17 x 10^-9^ |
| BAL217 | 1.96 ± 0.40 x 10^-9^ | N/A | 3.95 ± 1.18 x 10^-9^ | 8.08 ± 4.83 x 10^-9^ | 1.30 ± 9.16 x 10^-9^ | 1.23 ± 0.34 x 10^-7^ | 1.16 ± 0.51 x 10^-9^ | 4.77 ± 2.40 x 10^-10^ |
| BAL268 | N/A | N/A | 2.47 ± 0.71 x 10^-9^ | N/A | N/A | 6.96 ± 2.49 x 10^-9^ | N/A | N/A |
| BAL65 | N/A | 1.13 ± 0.19 x 10^-8^ | 8.79 ± 2.02 x 10^-9^ | 2.43 ± 1.77 x 10^-7^ | N/A | 1.56 ± 0.42 x 10^-8^ | 1.84 ± 0.49 x 10^-9^ | 7.96 ± 2.75 x 10^-10^ |
| BAL242 | N/A | 8.10 ± 1.22 x 10^-9^ | 8.02 ± 2.79 x 10^-9^ | N/A | N/A | 9.65 ± 3.45 x 10^-8^ | 1.05 ± 0.58 x 10^-9^ | 2.18 ± 1.20 x 10^-9^ |
| BAL179 | 1.67 ± 1.09 x 10^-9^ | 1.85 ± 0.23 x 10^-9^ | 1.52 ± 0.24 x 10-9 | 6.55 ± 3.91 x 10^-9^ | 9.87 ± 8.63 x 10^-9^ | 2.45 ± 0.42 x 10^-8^ | 2.02 ± 0.30 x 10^-9^ | 1.30 ± 0.20 x 10^-9^ |
| BAL248 | N/A | 3.36 ± 1.23 x 10^-8^ | 2.41 ± 0.66 x 10^-8^ | N/A | N/A | 2.06 ± 0.37 x 10^-7^ | 3.38 ± 1.35 x 10^-9^ | 7.08 ± 2.63 x 10^-9^ |
| BAL115 | N/A | 2.25 ± 0.42 x 10^-8^ | 4.18 ± 1.58 x 10^-9^ | N/A | N/A | 5.94 ± 2.20 x 10^-8^ | 2.33 ± 0.30 x 10^-8^ | 1.04 ± 0.32 x 10^-8^ |
| BAL213 | N/A | 2.50 ± 1.07 x 10^-8^ | 1.87 ± 0.63 x 10^-8^ | N/A | N/A | N/A | 3.93 ± 1.00 x 10^-8^ | 4.02 ± 1.23 x 10^-8^ |

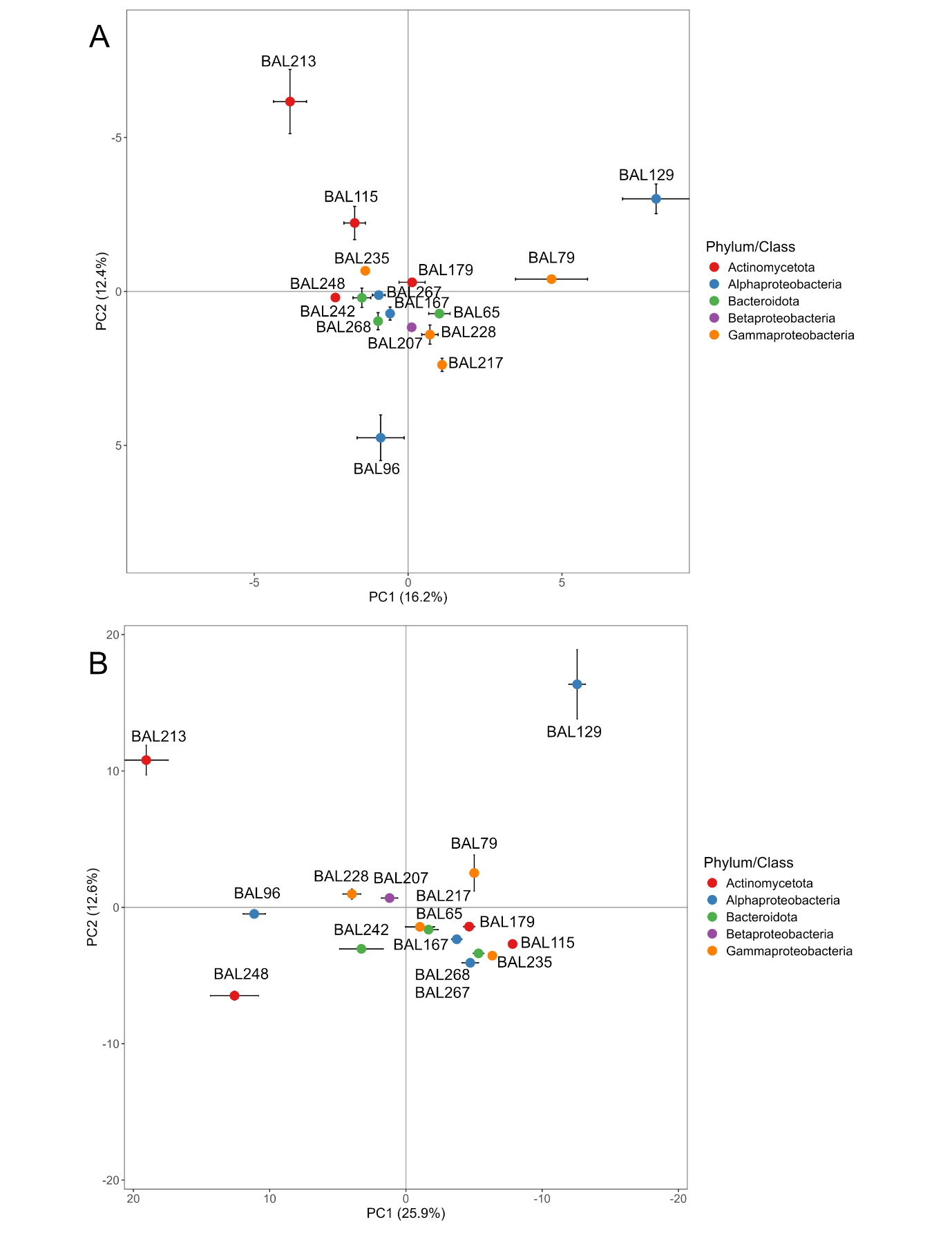

**Figure S1.** Principal component analysis (PCA) score plot of BVOC emissions for the 16 strains of A) 10 most abundant *m/z* per strain and B) all *m/z* except the 10 most abundant *m/z* (nmol cell^-1^ h^-1^, mean PC score position ± SE, n=3 except BAL79 = 2). The total volatilome is represented by 268 unique *m/z*.
